# Temporal kinetics of RNAemia and associated systemic cytokines in hospitalized COVID-19 patients

**DOI:** 10.1101/2020.12.17.423376

**Authors:** Debby van Riel, Carmen W.E. Embregts, Gregorius J. Sips, Johannes P.C. van den Akker, Henrik Endeman, Els van Nood, Jeroen van Kampen, Richard Molenkamp, Marion Koopmans, David van de Vijver, Corine H. GeurtsvanKessel

## Abstract

COVID-19 is associated to a wide range of extra-respiratory complications, of which the pathogenesis is currently not fully understood. In this study we report the temporal kinetics of viral RNA and inflammatory cytokines and chemokines in serum during the course of COVID-19. We show that a RNAemia occurs more frequently and lasts longer in patients that develop critical disease compared to patients that develop moderate or severe disease. Furthermore we show that concentrations of IL-10 and MCP-1—but not IL-6—are associated with viral load in serum. However, higher levels of IL-6 were associated with the development of critical disease. The direct association of inflammatory cytokines with viral load or disease severity highlights the complexity of systemic inflammatory response and the role of systemic viral spread.

## Brief report

COVID-19 has been associated with a wide range of extra-respiratory complications, including neurologic, cardiac, and thromboembolic complications. The pathogenesis of these extra-respiratory complications is not fully understood but several mechanisms are thought to contribute, including the systemic spread of SARS-CoV-2 and systemic inflammatory cytokines (Gupta et al., 2020). Even though SARS-CoV-2 viral RNA and inflammatory cytokines have been detected in the blood, their kinetics during infection are poorly understood and it is unclear if either RNAemia or inflammatory cytokines are associated with each other or with other disease parameters. We hypothesize that RNAemia occurs frequently during severe COVID-19 and that it contributes to the systemic cytokine responses. Therefore we aimed to get insight in the kinetics of SARS CoV-2 RNAemia and the associated systemic inflammatory response.

Diagnostic specimen of twenty patients (16 male, 4 female), hospitalized at the Erasmus MC in the Netherlands in March and April 2020, were analyzed. Ten patients developed moderate or severe disease (grouped together) and 10 critical disease, according to NIH disease severity guidelines (https://www.covid19treatmentguidelines.nih.gov/overview/management-of-covid-19). One or more underlying diseases were present in 19 patients which included cardiovascular diseases (11), diabetes (9), previous cerebro-vascular accident (4), previous respiratory disease (3), cancer (3), solid organ transplantation (2), kidney failure (1), psychosis (1) or sarcoidosis (1). None of the patients received dexamesthasone during the course of disease. Further patient characteristics are included in Supplemental Table 1. A total of 176 serum samples were analyzed and a paired RT ([naso]pharyngeal swab or sputum) was available for 131 specimens (Table S2).

**Table 1:** Concentrations of individual cytokines and associations with RNAemia, a ct value in respiratory samples <30, development of critical disease, fatal disease outcome and the first ten days after disease onset. Values with the group names indicate the number of patients and the number of samples (patients [number of samples]) within the specified group. Univariate generalized estimated equations were performed on the individual cytokines (log10 transformed) and associations with a p <0.1 were included in a multivariate analysis.

| Association with RNAemia |  |  |  |  |
| --- | --- | --- | --- | --- |
|  | Median concentration pg/mL (range) |  | P- value |  |
|  | Ct <45 in serum (13 [58]) | Ct =45 in serum (19 [72]) | Univariate | Multivariate |
| IFNγ | 3.0 (3-385) | 3.0 (3-138) | **0.003** | 0.528 |
| IL1β | 2.0 (0.8-4.8) | 0.8 (0.8-3.5) | **0.046** | 0.099 |
| IL-2 | 4.2 (2.1-13.9) | 2.1 (2.1-31.1) | **0.059** | 0.969 |
| IL-6 | 203 (10-4090) | 61 (8-1055) | 0.018 | 0.103 |
| IL-8 | 25.4 (4.6-99.8) | 11.7 (3.4-57.5) | **<0.001** | 0.251 |
| IL-10 | 11.5 (1.7-140) | 1.7 (1.7-26.2) | **<0.001** | **<0.001** |
| IL-17A | 3.2 (3.2-13.3) | 3.2 (3.2-19.4) | 0.150 | - |
| IP-10 | 3683 (168-9155) | 1380 (86-7282) | **<0.001** | 0.170 |
| MCP-1 | 272 (73-2500) | 184 (70-2500) | **0.019** | **0.036** |
| TNFα | 1.2 (1.2-10.7) | 1.2 (1.2-9.8) | 0.410 | - |

| Association with a high viral load(ct <30) in respiratory tract |  |  |  |  |
| --- | --- | --- | --- | --- |
|  | Median concentration pg/mL (range) |  | P- value |  |
| | Ct <30 in RT (18 [82]) | Ct ≥30 in RT (15 [32]) | Univariate | Multivariate |
| IFNγ | 3.0 (3-385) | 3.0 (3-11.5) | **0.007** | **0.015** |
| IL1β | 1.8 (0.8-4.8) | 0.8 (0.8-2.8) | **0.008** | **<0.001** |
| IL-2 | 3.8 (2.1-13.9) | 2.1 (2.1-16.1) | **0.014** | 0.686 |
| IL-6 | 143 (10-4090) | 82 (8-768) | **0.029** | 0.749 |
| IL-8 | 20 (4.4-99.8) | 11.7 (3.4-40.8) | **0.002** | 0.269 |
| IL-10 | 6.4 (1.7-141) | 1.7 (1.7-9.7) | **0.003** | 0.271 |
| IL-17A | 3.2 (3.2-19.4) | 3.2 (3.2-13.8) | 0.670 | - |
| IP-10 | 2686 (86-9155) | 1123 (123-6656) | **0.021** | 0.759 |
| MCP-1 | 239 (79-2500) | 154 (73-2500) | **<0.001** | 0.358 |
| TNFα | 1.2 (1.2-9.8) | 1.2 (1.12-10.7) | 0.823 | - |

| Association with the development of critical disease |  |  |  |  |
| --- | --- | --- | --- | --- |
|  | Median concentration pg/mL (range) |  | P- value |  |
|  | Critical disease (10 [91]) | Moderate or severe disease (10 [39]) | Univariate | Multivariate |
| IFNγ | 3.0 (3-73) | 6.0 (3-385) | 0.084 | 0.228 |
| IL1β | 2.1 (0.8-4.8) | 0.8 (0.8-2.0) | **0.003** | **0.003** |
| IL-2 | 3.6 (2.1-13.9) | 2.2 (2.1-31.1) | 0.440 | - |
| IL-6 | 144 (10-4090) | 49.8 (8.1-254) | **0.041** | **0.005** |
| IL-8 | 19.4 (3.4-99.8) | 12.4 (14.5-29.9) | 0.140 | - |
| IL-10 | 5.7 (1.7-141) | 5.7 (1.7-67.9) | 0.620 | - |
| IL-17A | 3.2 (3.2-19.4) | 3.2 (3.2-14.9) | 0.700 | - |
| IP-10 | 1918 (86-9155) | 2487 (692-8015) | 0.121 | - |
| MCP-1 | 197 (73-2500) | 316 (70-2500) | 0.480 | - |
| TNFα | 1.2 (1.2-10.7) | 1.2 (1.2-9.7) | 0.373 | - |

| Association with fatal outcome |  |  |  |  |
| --- | --- | --- | --- | --- |
|  | Median concentration pg/mL (range) |  | P-value |  |
|  | Survivor (15 [80]) | Non-survivor (5 [50]) | Univariate | Multivariate |
| IFNγ | 3.0 (3-385) | 3 (3-73) | 0.880 | - |
| IL1β | 0.8 (0.8-3.5) | 1.9 (0.8-4.8) | 0.157 | - |
| IL-2 | 2.3 (2.1-31.1) | 4.0 (2.1-13.9) | 0.149 | - |
| IL-6 | 56 (8-4090) | 270 (10-3930) | **0.031** | 0.063 |
| IL-8 | 13.4 (3.4-60.9) | 23.9 (4.6-99.8) | 0.070 | 0.607 |
| IL-10 | 3.3 (1.7-140.8) | 6.6 (1.7-67.9) | 0.610 | - |
| IL-17A | 3.2 (3.2-14.9) | 3.2 (3.2-19.4) | 0.210 | - |
| IP-10 | 1922 (86-8015) | 2601 (168-9155) | 0.160 | - |
| MCP-1 | 217 (70-2500) | 247 (73-2500) | 0.470 | - |
| TNFα | 1.2 (1.2-9.7) | 1.2 (1.2-10.7) | 0.930 | - |

| Association with the first ten days post disease onset |  |  |  |  |
| --- | --- | --- | --- | --- |
|  | Median concentration pg/mL (range) |  | P-value |  |
| | ≤ 10 days (14 [51]) | >10 days (18 [79]) | Univariate | Multivariate |
| IFNγ | 6 (3-385) | 3 (3-31.1) | **<0.001** | **0.034** |
| IL1β | 1.3 (0.8-4.8) | 1.8 (0.8-3.88) | 0.330 | - |
| IL-2 | 3.1 (2.1-31.1) | 3.5 (2.1-16.1) | 0.690 | - |
| IL-6 | 144 (10-3930) | 74 (8-4090) | 0.147 | - |
| IL-8 | 17 (4.8-99.8) | 14.7 (3.4-60.9) | 0.360 | - |
| IL-10 | 7 (1.7-140.8) | 1.8 (1.7-131.9) | 0.182 | - |
| IL-17A | 3.2 (3.2-6.1) | 3.2 (3.2-19.4) | 0.400 | - |
| IP-10 | 3683 (252-9155) | 1653 (86-7646) | **0.013** | 0.058 |
| MCP-1 | 320 (70-2500) | 170 (73-2500) | 0.190 | - |
| TNFα | 1.2 (1.2-9.8) | 1.2 (1.2-10.7) | 0.800 | - |

Viral RNA was measured by qPCR for the E gene in all serum and RT samples as described previously (Corman et al., 2020). RNAemia was detected in 50% and 90% of the patients with respectively moderate/severe or critical disease. The duration of the RNAemia per patient could not be determined, since samples early after disease onset were not available for all patients, but RNAemia was more frequently detected from 11 days post disease onset (dpd) in patients who developed critical disease (OR 4.65 CI 1.05-20.64; p = 0.038; Figure 1A, Table S2), as was calculated using generalized estimated equations, corrected for repeated samples within patients. Furthermore, independent of disease severity, RNAemia was associated with a ct value <30 in paired RT samples (OR 9.47, CI 3.08-29.07; p=0.009) and < 11 dpd (OR 4.61, CI 1.80-11.87; p=0.001; figure 1B). In this study there was no correlation between RNAemia and age (OR 1.0, CI 0.99-1.01; p=0.31) or BMI (OR1.0, CI 0.9-1.2; p=0.58). Virus could not be isolated from serum samples, but since these had been collected for molecular and serological diagnostics, the sample handling and storage was most likely suboptimal for virus culture. In addition, we have previously shown that virus is difficult to culture from samples with an ct value of 27 (van Kampen J.J.A, 2020).

**Figure 1:**
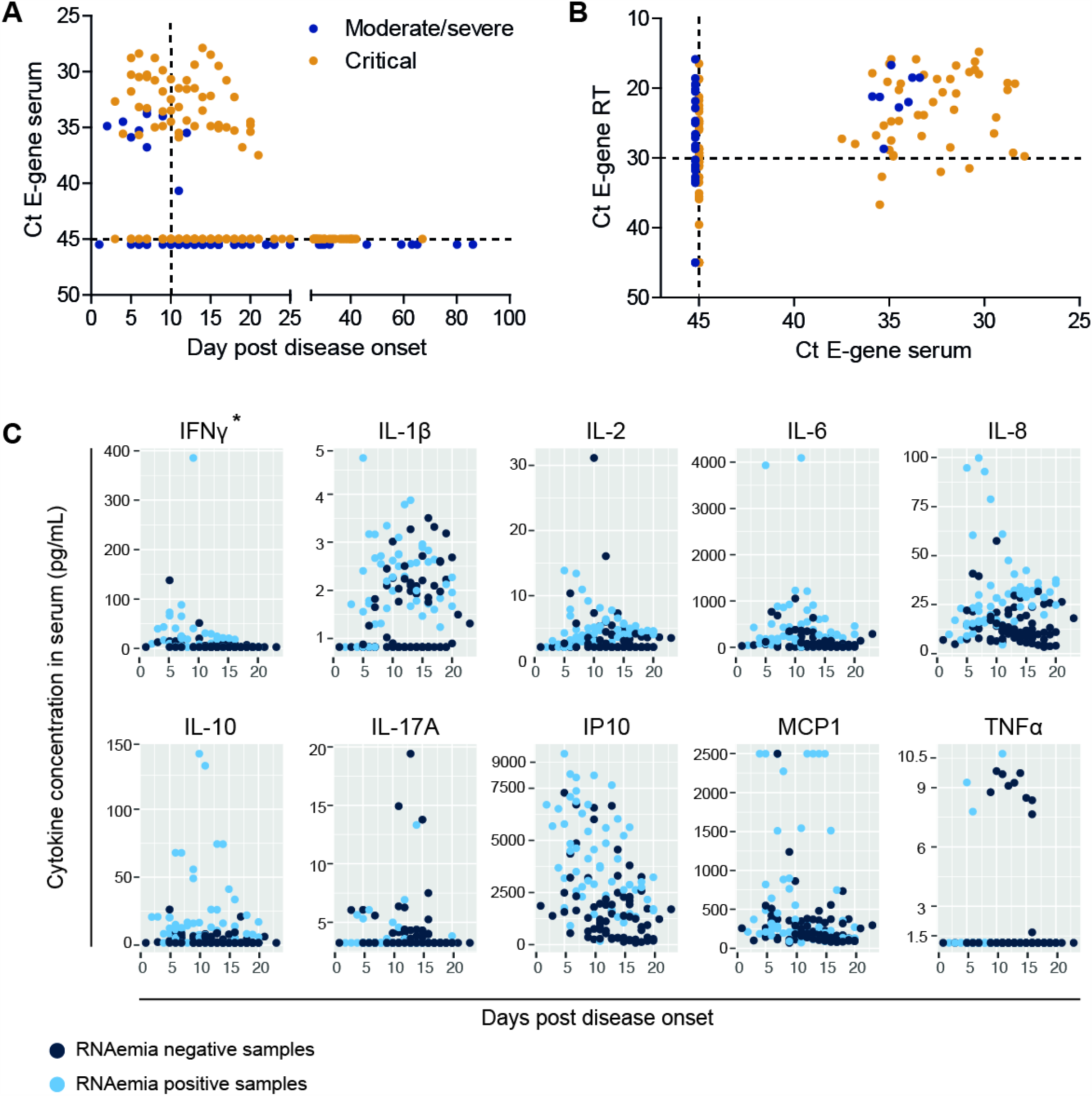
RNAemia and systemic inflammatory responses in hospitalized COVID-19 patients. The detection of SARS-CoV-2 viral RNA (E-gene) in serum in patients with moderate/severe or critical disease plotted against the day post disease onset (A). The detection of SARS-CoV-2 viral RNA in serum plotted against the detection of SARS-CoV-2 viral RNA in respiratory sample. The dashed line represents a ct value of 30 in respiratory samples (B). Individual cytokines plotted against the day post disease onset. * refers to significant higher levels in multivariate analyses < 10 dpd compared to > 10 dpd (C).

Next, we used a 13-plex cytometric bead assay to determine kinetics of the inflammatory cytokines and chemokines IL-1β, IL-2, IL-4, IL-6, IL-8, IL-10, IL-12p70, IL-17A, IP-10, IFNγ, TNFα, MCP-1 and TGFβ in serum of patients up to 21 dpd (Fig S1). All tested cytokines were significantly upregulated during the course of infection, with the exception of IL-4, IL-12p70 and TGFβ, which were therefore excluded from further analysis. IFNγ, IL-1β, IL-2, IL-6, IL-8, IL-10, IL-17A, IP-10, MCP-1 and TNFα were induced during the course of disease (Figure 1C), although large differences were observed among individual patients. Overall, a dynamic response was observed in the majority of patients (Figure S1). Generalized estimated equations, corrected for repeated sampling within patients, were used to determine associations of individual cytokines with RNAemia, a ct RT <30, disease severity and outcome, and period after disease onset. Of all cytokines and chemokines only IFNγ was associated with a disease period of ≤10 days in both uni- and multivariate analysis (Table, Figure 1C).

First, we determined which cytokines or chemokines were associated with the viral load in serum or RT sample. A RNAemia was associated with increased levels of IL-1β, IL-6, IL-8, IL-10, IP-10 and MCP-1 in a univariate analysis. Subsequent multivariate analyses, adjusting for the ct value in the RT, showed that Il-10 and MCP-1 were independently associated with RNAemia (Table, Figure S2A). In a univariate analysis, ct value <30 in the RT was associated with increased serum levels of IFNγ, IL-1β, IL-2, IL-6, IL-8, IL-10, IP-10 and MCP-1 compared to a ct value >31 in the RT, and with increased levels of IFNγ and IL-1β in a multivariate analysis (Table, Figure S2B).

Next, we determined which cytokines or chemokines were associated with the development of critical disease and death as outcome. Patients that developed critical disease had significantly higher levels of IL-1β and IL-6 compared to patients that developed moderate or severe disease (Table, Figure S3A). Death as an outcome was not associated with elevated levels of any of the cytokine included in this study in the multivariate analyses (Table, Figure S3B).

Altogether, this study shows the presence of RNAemia in critically ill COVID-19 patients. The overall detection of RNAemia in 70% of patients included in this study is higher than in most other reports (Hogan et al., 2020, Fajnzylber et al., 2020). This is likely due to the fact that all patients included in this study were hospitalized with moderate, severe or critical disease and the fact that we analyzed samples throughout the course of disease. The fact that a RNAemia is detected at later time points post disease onset in patients that develop critical disease, together with the detection of SARS-CoV-2 in extra-respiratory tissues (Gupta et al., 2020), suggests the systemic spread of SARS-CoV-2 contributes to the pathogenesis of severe COVID-19.

The role of systemic cytokines during the course of COVID-19 is only partly understood. In line with previous studies we show that higher levels of IL-6 are associated with the development of more severe disease (Del Valle et al., 2020, Zhang et al., 2020). However, IL-6 is not directly associated with viral load in either serum or the respiratory tract in our study, which suggest that IL-6 production is not directly triggered by virus replication. In contrast, IL-10, MCP-1, IL-1β and IFNγ were associated with either viral RNA in serum or RT suggesting that these were triggered—either directly or indirectly—by virus replication. This suggests that the systemic cytokine response is complex, but is at least in part directly associated with virus within the respiratory tract or circulation. The multifaceted mechanism of systemic responses during the course of disease and the exact role of these systemic responses in the pathogenesis of COVID-19 requires further characterization.

Even though this study has several limitations, such as the number of patients, the retrospective character and the usage of diagnostic samples, it reveals important insight on the temporal kinetics of both RNAemia and associated systemic cytokine responses. Our findings suggest that both RNAemia and systemic responses contribute—at least in part—to the systemic pathogenesis of severe COVID-19, which fits with the wide spectrum of extra-respiratory complications associated with COVID-19. Furthermore, we show that in order to acquire more insights into the systemic pathogenesis, it is essential to analyse the kinetics of systemic viral load and responses during the course of disease. This knowledge will be essential for the development and timing of intervention strategies that target either the host immune response or virus replication

## Acknowledgments

The authors gratefully acknowledge Matthijs Raadsen for analytical support. Debby van Riel is supported by the Netherlands Organization for Scientific Research (VIDI 91718308) and a EUR fellowship. In addition, this work supported by funding from the European Union’s Horizon 2020 research and innovation program under grant agreement No. 101003589 (RECoVER)

## Supplemental materials& methods, figures and tables

### Materials and methods

#### Patient specimen and data collection

Diagnostic respiratory and serum samples of COVID-19 patients admitted to Erasmus are sent to the unit of clinical virology, Viroscience department, Erasmus MC. In 20 patients admitted in March and April 2020, we performed qPCRs on serum samples collected for diagnostic purposes during admission, as previously described (ref van Kampen Nat Comm). Other available diagnostic results (serology, virus culture and qPCR on respiratory tract specimen) were extracted from our diagnostic laboratory information management system. In addition, following information was extracted from the electronic patient files: date of onset of symptoms, disease severity (hospitalized on ICU with mechanical ventilation, hospitalized on ICU with oxygen therapy, hospitalized to ward with oxygen therapy, hospitalized to ward without oxygen therapy), whether the patients were still alive or not when they were discharged.

#### Medical ethical approval

All patient specimen and data used in this study were collected in the context of routine clinical patient care. Additional analyses were performed only on surplus of patient material collected in the context of routine clinical patient care. Our institutional review board approved the use of these data and samples (METC-2015-306). METC-2015-306 is a generic protocol to study viral diseases. Informed consent was waived by the privacy knowledge office of the Erasmus MC.

#### Cytometric bead assay

Systemic cytokines were quantified in all patients’ sera up to 21 dpd. Sera were analysed in duplo using the 13-plex Human essential immune Legendplex panel (Biolegend). A standard cytokine cocktail was taken along in duplo to allow for quantification of the cytokine concentrations. The cytometric bead assay was performed on a FACSLyric (BD) and the data were analysed using the Legendplex data analysis software.

#### Analyses

Statistical analysis was performed using R version 4.0.3 and the geepack package. Associations between presence of RNAemia and a Ct <30 in respiratory samples and a dpd <11 were determined after dichotomizing the values and evaluating the fit according to the quasi-likelihood under the independence criterion (QIC) by generalized estimating equations corrected for multiple samples per patient. Associations between individual cytokines and RNAemia, a ct <30 in respiratory samples, critical disease and a fatal outcome were assessed using generalized estimating equations corrected for multiple samples per patient. Associations of cytokines with a p-value of <0.1 were subjected to a multivariate analysis.

**Figure S1:**
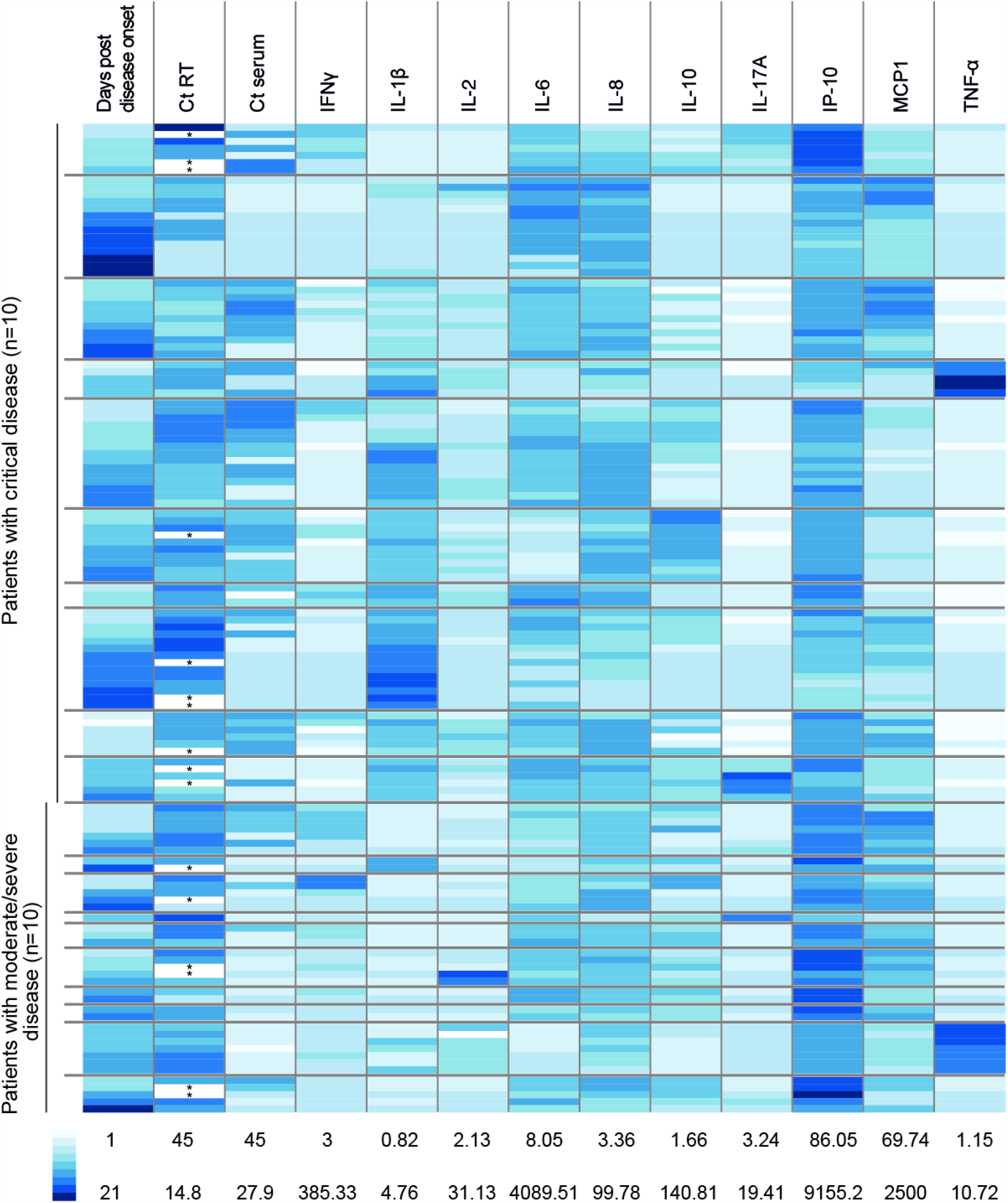
Individual patient data of days post disease onset, ct value in the respiratory tract (RT), ct value in serum and the concentration of respective cytokines in pg/mL.

**Figure S2:**
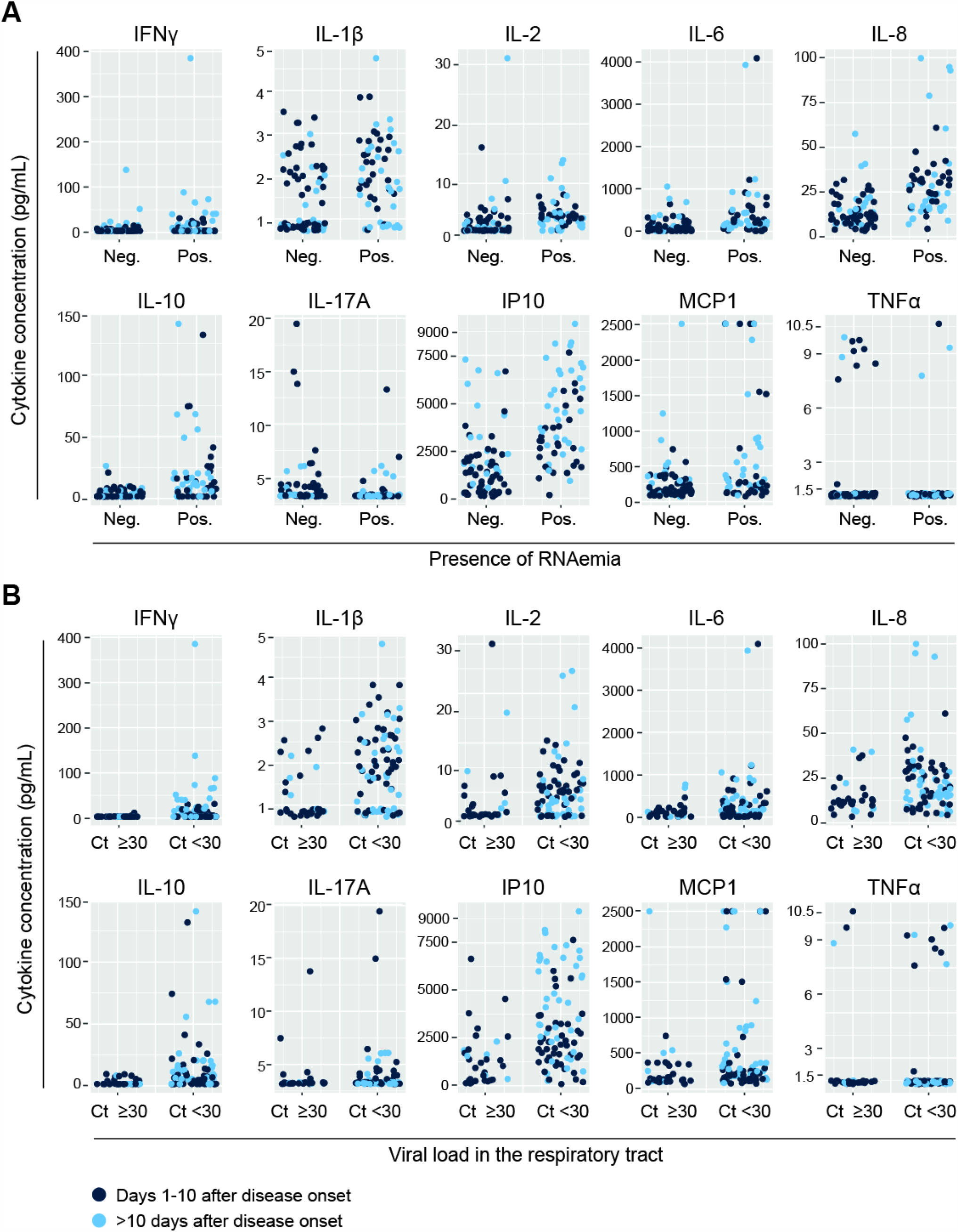
Scatterplots of all individual cytokine or chemokine measurements, plotted against the presence of RNAemia in serum (A) or a ct in the respiratory tract <30 (B). Samples are divided between 1-10 dpd (light blue) and >10 dpd (dark blue).

**Figure S3:**
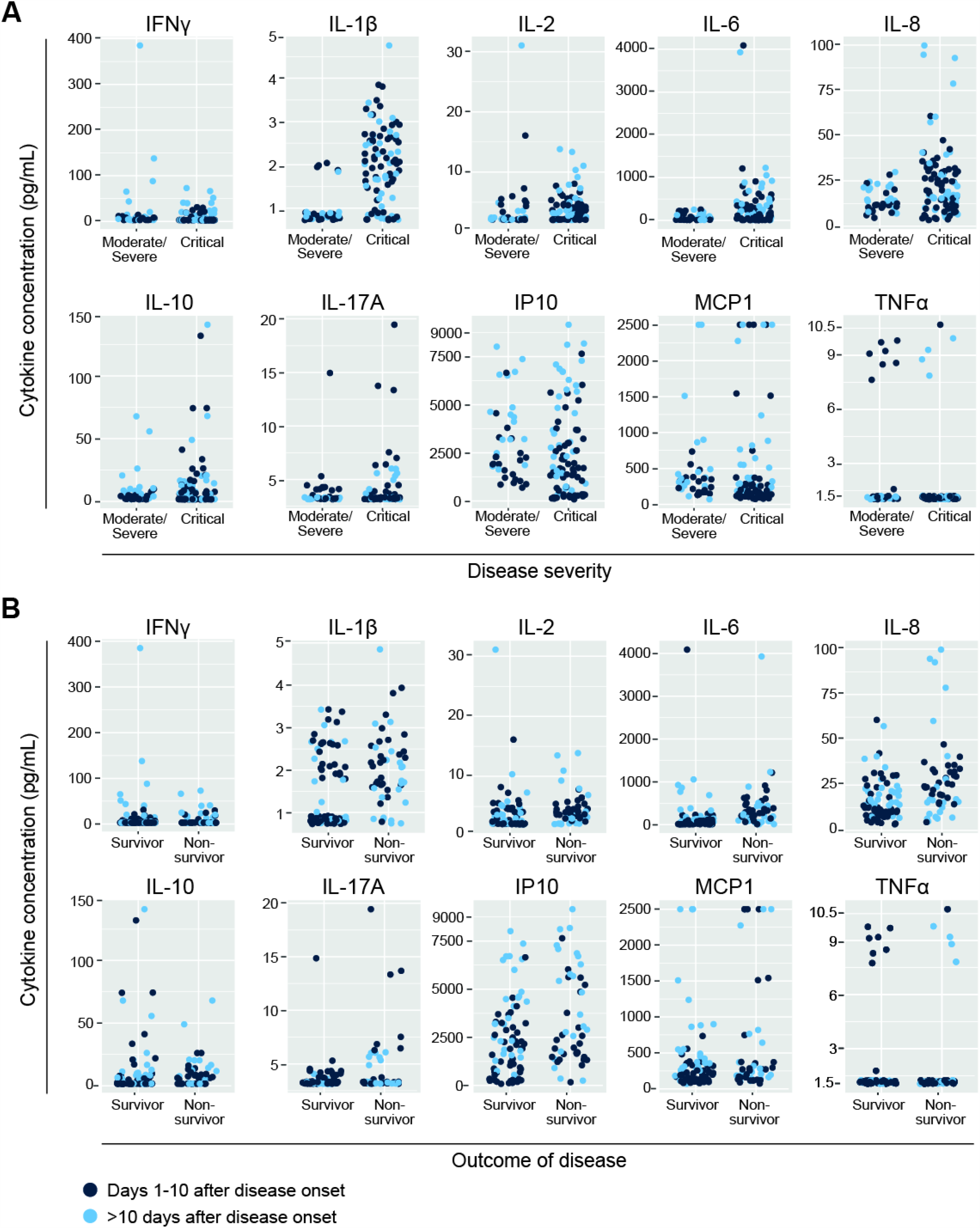
Scatterplots of all individual cytokine of chemokine measurements, plotted against the severity of disease (A) or the outcome of disease (B). Samples are divided between 1-10 dpd (light blue) and >10 dpd (dark blue).

**Table S1:** Information of patients included in the study

|  |  | Moderate/Severe disease | Critical disease | Total |
| --- | --- | --- | --- | --- |
| **Total** |  | 10 | 10 | 20 |
| **Gender** | **Male** | 9 | 7 | 16 |
|  | **Female** | 1 | 3 | 4 |
| **Age\*** | **Mean** | 60.8 | 59.3 | 60.1 |
|  | **Range** | 25-84 | 17-84 | 17-84 |
| **BMI\*** | **Mean** | 26.3 | 29.2 | 28 |
|  | **Range** | 19-33 | 24-38 | 19-38 |
| **Outcome** | **Death** | 0 | 5 | 5 |

\* no significant differences between patients with moderate/severe and critical disease.

**Table S2:** Detection of viral RNA serum of patients with moderate/severe or critical disease. Analyses are done on the total number of samples, and on samples from 1-10 days post disease onset (dpd) and >10 dpd.

|  |  | Patient with moderate or severe disease (samples) | Patient with critical disease (samples) | Total number of patients (samples) |
| --- | --- | --- | --- | --- |
| **Number of samples** | Total | 10 (57) | 10 (119) | 20 (176) |
|  | 1-10 dpd | 6 (17) | 8 (35) | 14 (52) |
|  | >10 dpd | 10 (40) | 9 (84) | 19 (124) |
| **PCR positive samples (E-gene)** | Total | 5 (10) | 9 (51) | 14 (61) |
|  | 1-10 dpd | 4 (8) | 7 (23) | 11 (31) |
|  | >10 dpd* | 2 (2) | 7 (28) | 9 (30) |
| **% positive samples** | Total | 50 (18) | 90 (43) | 70 (35) |
|  | 1-10 dpd | 67 (47) | 88 (66) | 79 (60) |
|  | >10 dpd | 20 (5) | 78 (33) | 47 (24) |
| **RNAemia** | Mean spd** | 7 | 11.6 | 10.7 |
|  | Range dpd | 2-12 | 4-21 | 2-21 |

\* p<0.05 between patients with moderate/severe and critical disease
\*\* p<0.01 between patients with moderate/severe and critical disease

